# ESPERDYNE: A Dual-Band Heterodyne Monitor and Ultrasound Recorder for Bioacoustic Field Surveys

**DOI:** 10.1101/2025.08.30.673206

**Authors:** Ravi Umadi

## Abstract

**Background:** Ultrasonic monitoring is essential for ecological studies of bats and other animals, yet high-performance field devices remain prohibitively expensive and inaccessible–particularly in biodiversity-rich regions with limited research infrastructure. Existing low-cost options often lack real-time listening and recording features. There remains a critical need for versatile, affordable, and field-ready tools that support acoustic behavioural research, educational and conservation outreach.

**New Tool:** I introduce *Esperdyne*, an open-source, dual-channel ultrasound monitoring and recording system based on the ESP32-S3 microcontroller. With a component cost under €75, Esperdyne combines real-time heterodyne monitoring, stereo recording from a retroactive ring buffer, and an intuitive rotary-based user interface with OLED display. It supports full-duplex 192 kHz audio, dual-band tuning for simultaneous FM/CF monitoring, and real-time playback via headphones or a speaker. All audio processing–including adjustable carrier frequency mixing, gain control, and file-saving logic–is implemented without reliance on fixed-rate audio libraries.

**Applications:** Esperdyne has been tested in field conditions and shown to reliably detect high-SNR calls and harmonics from free-flying bats. A companion tool *Bat Reviewer* supports rapid inspection, playback, and export of selected recordings. Together, these tools enable portable, solo-operated acoustic surveys with minimal training. Beyond ecological research, Esperdyne is suitable for education, outreach, and preliminary field assessments in remote or resource-constrained settings. Its modular design encourages hardware customisation and firmware extension by interdisciplinary teams.

**Availability and Implementation:** Full hardware schematics, firmware, and software tools are publicly available. The system can be built using hobbyist-accessible components and standard Arduino tooling. By sharing this system openly, I aim to lower technical barriers and foster broader participation in ultrasound-based biodiversitymonitoring and conservation. Esperdyne demonstrates how microcontroller-based platforms can bridge gaps between affordability, usability, and scientific capability–supporting global efforts in soundscape ecology.

## 1 INTRODUCTION

Acoustic monitoring of bat calls is a widely used method for species surveys, identification, and even the discovery of previously unknown taxa [1–4]. It is also a key educational tool in sound-scape ecology, providing an accessible gateway into understanding animal communication and nocturnal biodiversity [5–10]. The most effective tools for active field surveys—particularly when tracking bats, on the move through challenging terrain—are lightweight, reliable, and intuitive to operate [11].

Despite recent advances, the range of available heterodyne detectors and field recorders remains limited, especially in the lower cost segment. Many professional-grade devices cost several hundred to several thousand euros. For example, one of the most widely used heterodyne-enabled field recorders, the Pettersson D1000X[12], retails for approximately €3600, weighs 600 grams (excluding batteries), and requires CompactFlash cards for storage. Other commercial bat detectors with recording capabilities are typically priced around €1400, while non-recording devices often start at €250. In Germany, the SSF BAT3-Detektor is a commonly used monitoring unit with limited recording capabilities and a cost of approximately €350. Handheld devices from manufacturers like Dodotronics provide mobile-compatible plug-and-play solutions, but often require additional proprietary software and cost over €250.

While these devices offer robust features and professional reliability, their high cost remains a significant barrier—particularly for educators, students, and conservation projects in the Global South. From personal experience, I recall the considerable challenge of acquiring a bat detector early in my research career in India: the device cost over a month’s salary and incurred additional import duties. I was struck by the irony that regions rich in bat diversity often lack access to the tools necessary to study them. In contrast, the tools originate from areas with relatively low bat diversity.

Open-source projects such as AudioMoth [13] and RPi Bat Projekt [14] have addressed part of this gap by providing low-power, autonomous passive recording platforms. These systems are well-suited for long-term monitoring and programmable deployment but lack the capability for real-time listening and user-directed recording, which can be critical for behavioural studies and educational outreach.

I recently developed a multichannel field recorder for bioacoustic behavioural experiments as part of my ongoing work in *embedded ultrasonics*. Combined with custom microphone arrays built using a modular toolkit [15–17], this system offers both real-time listening and retroactive capture of ultrasonic events, enabling novel experimental paradigms. Building on this foundation, I present the design and implementation of *Esperdyne*^1^—a compact, low-cost, open-source dual-band heterodyne monitor and stereo ultrasound recorder built around the ESP32-S3 microcontroller.

Esperdyne allows simultaneous tuning of two frequency bands, enabling live parallel monitoring of frequency-modulated (FM) and constant-frequency (CF) bats—a feature rarely found even in high-end detectors. The system features an intuitive user interface with rotary-encoder-based control, OLED display, real-time headphone and speaker output, and a tap-to-save ring buffer allowing up to 20 seconds of retroactive recording. It is built entirely from off-the-shelf, hobbyist-accessible components, with fully open-source firmware and hardware.

The prototype was assembled on a perforated PCBand field-tested during the summer of 2025 along the river Isar in southern Germany. Figures in this paper provide representative recordings, and reference files from these surveys are available for comparison and testing by users building their own units.

The Esperdyne provided a high-fidelity listening experience with minimal self-noise—an issue common in other bat detectors that artificially boost gain to reveal weak signals. The perceived audio quality depends strongly on the microphone and DAC components used, yet the system, as tested, produced results comparable to commercial units costing over 10 times as much. With a build cost under €75, Esperdyne offers a powerful, affordable alternative for researchers, educators, and students worldwide.

Complementing the hardware, I also developed *Bat Reviewer*, a lightweight MATLAB-based graphical interface forbrowsing, inspecting, and curating ultrasonic recordings – available as cross-platform standalone executables [18]. The tool enables users to navigate entire recording sessions, visualise waveforms and spectrograms, and listen to heterodyned audio in real time with adjustable carrier frequency and playback volume. Export functions streamline the selection of high-quality calls and facilitate organised data curation. Combining visual inspection with auditory evaluation in a single interface, *Bat Reviewer* shortens the review cycle and supports more systematic field data processing.

In summary, Esperdyne and BatReviewer demonstrate how open-source, low-cost solutions can lower barriers to acoustic monitoring. By coupling accessible hardware with flexible software, these tools provide a practical foundation for research and educational applications, particularly in regions where commercial systems are prohibitively expensive. Beyond immediate utility, they also highlight the potential of community-driven development in advancing the field of bioacoustics and ensuring broader participation in biodiversity monitoring.

## 2 METHODS

The Esperdyne system is built around the ESP32-S3 (Espressif Systems (Shanghai) Co., Ltd.) [19] microcontroller, selected for its dual-core architecture, onboard Pseudo-Static Random Access Memory (PSRAM), and two independent I2S ports—crucial for full-duplex ultrasonic audio processing. It integrates a high-fidelity WM8782 24-bit stereo ADC (Cirrus Logic, Inc., Austin, Texas, USA) [20] for input and a PCM5102A DAC (Texas Instruments, Dallas, Texas, USA) [21] for audio output.

Audio from the SPU0410LR5H-QB analogue omnidirectional MEMS ultrasonic microphones (Knowles Electronics, LLC. Itasca, IL, USA) is digitised via the WM8782 ADCand captured through the I2S0 input interface. The digitised samples are processed for heterodyne conversion (based on user-set carrier frequencies) and output via the PCM5102A DAC through I2S1. Audio is simultaneously written into a ring buffer stored in PSRAM, allowing buffering up to 20 seconds on the onboard 8MB PSRAM with a single channel at a given sampling rate and bit depth.

All components were assembled on 6×8 cm perforated prototyping PCBs using hand-soldered point-to-point wiring.

### 2.1 User Interface and OLED Display

The Esperdyne device features a minimal yet intuitive user interface built around a rotary encoder, three tactile buttons, and a 128*×*32 pixel OLED display. This interface allows real-time adjustment of heterodyne parameters and clear feedback on device status.

#### Rotary Encoder and Parameter Selection

The rotary encoder serves as a dual-mode control for adjusting the heterodyne **carrier frequency** (F) and **playback gain** (V). A short press on the encoder toggles between these two editable parameters. The currently selected parameter is indicated by displaying its corresponding label (F or V) on the OLED. Rotating the encoder clockwise increases the value, while counter-clockwise rotation decreases it. Adjustments are applied in real-time to the active audio channel.

#### Tap-to-Save

A separate *tap-to-save* button triggers the recording function, writing the last 5 seconds of buffered audio to the SD card as a WAV file. While writing the file, a message “Saving File” is displayed.

#### Channel 2 Frequency Adjustment

When the device is in frequency tuning mode, a dedicated button enables editing of the second heterodyne channel (CH2). By holding this button while rotating the encoder, users can independently adjust the carrier frequency for CH2, allowing both channels to be tuned separately for dual-band monitoring.

#### Listening Mode

A push button toggles the dual-channel heterodyned output between two modes. In *MX mode*, signals from both microphones are mixed and played back as a combined output on both channels. True stereo monitoring is enabled in *ST mode*, routing each microphone input to its corresponding output channel.

#### Display Toggle

An additional *screen toggle* button turns the OLED display on or off, allowing users to reduce power consumption during extended operation or eliminate screen glare in low-light environments.

#### OLED Layout and Visual Feedback

The OLED interface updates dynamically in response to user input, providing clear visual feedback. It uses the Adafruit_GFX library with FreeSerifand FreeSerifItalic fonts for enhanced legibility. The screen layout is structured as follows:

- **Top row:** Labels and units for all displayed parameters
- **Middle row:** Channel 1 frequency (left), current adjustment mode (F: Frequency / V: Volume), file index, and Channel 2 frequency (right)
- **Bottom row:** Output mode (MX/ST) on the left and device logo in the centre.

This layout ensures immediate visibility of all critical parameters, enabling quick and intuitive adjustments during field deployment.

### 2.2 Firmware Overview

The Esperdyne firmware is implemented using the Arduino framework. The operating system manages high-speed real-time audio input/output, heterodyne demodulation, and user interaction, all optimised for responsive field use. Below, I summarise the core architecture and signal flow.

#### 2.2.1 Signal Flow and I2S Configuration

Dual I2S peripherals are initialised for independent high-fidelity audio input and output. I2S0 is configured as the receiver, acquiring audio from external ADCs at 192 kHz with two channels. I2S1 acts as the transmitter, delivering processed audio to the DAC. The implementation logic may be found in supplementary material B.1.

#### 2.2.2 Ring Buffer Management

A 5 s stereo ring buffer is dynamically allocated in external PSRAM to store the most recent audio samples. This enables a *tap-to-save* feature, writing the last 5 seconds of audio to a WAV file on microSD upon user command. Buffer writes occur in real time in the main audio loop. The implementation logic can be found in the supplementary material B.2.

#### 2.2.3 User Interaction and Editable Parameters

The rotary encoder and push buttons provide intuitive control over frequency, volume, display state, recording action, and monitoring mode. Depending on input state, the encoder adjusts either the *frequency* or the *volume*, while the *channel 2 edit* button allows independent tuning. The implementation logic is provided in supplementary material B.3

#### 2.2.4 Saving and Playback

When the record button is pressed, the firmware stops I2S input, copies the ring buffer to a WAV file using a custom header writer, then resumes streaming. File names auto-increment to prevent overwrites (REC000.WAV to REC999.WAV).

The firmware prioritises low-latency response and stability, with state-based logic ensuring no recordings are interrupted mid-stream and all inputs are debounce-guarded. Delays are added strategically (e.g., post PSRAM allocation) to avoid race conditions during startup.

A splash screen and OLED UI provide visual feedback, showing frequencies, gain, mode, file index, and playback mode in a compact 128×32 layout (see Figure 1).

**Figure 1:**
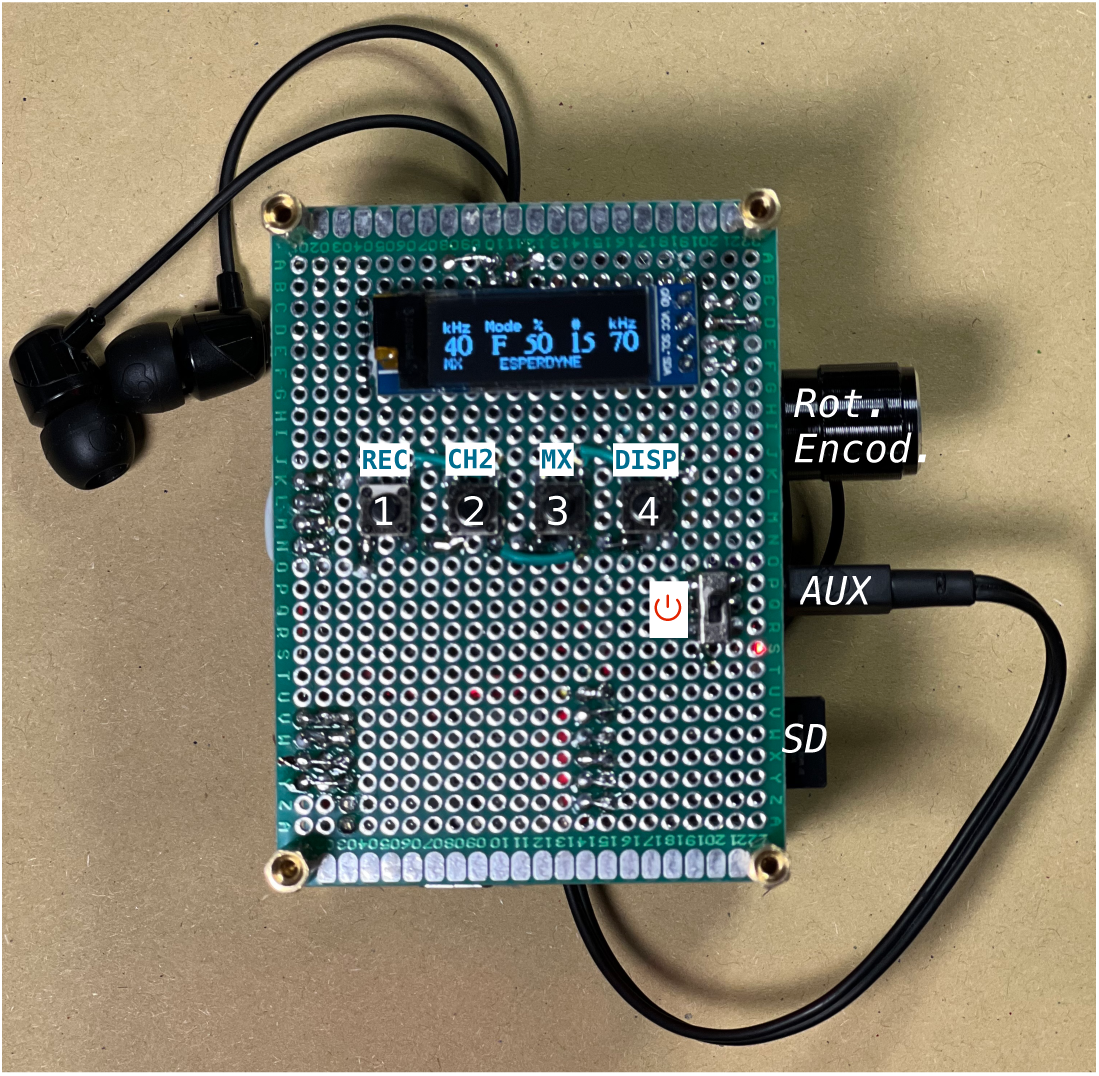
Esperdyne Field Recorder. The device is based on the ESP32-S3 MCU with dual I2S input, rotary encoder, OLED display, and microSD card. It includes a tap-to-save recording button (REC), channel 2 tuning hold button (CH2), and an OLED screen toggle (DISP). The front face includes ports and controls: SD card slot, rotary encoder, AUX Output, and power button. The interface allows real-time stereo heterodyne output with independent frequency tuning for each channel, while a ring buffer stores the last 5 seconds of audio, triggered for saving upon user input. The two carrier frequencies are tuned to 40 kHz and 70 kHz, and the display shows the current number of recorded files to be 15. The volume is set at 50 %.

### 2.3 I^2^S Configuration

Both I2S0 and I2S1 are set to operate at a 192 kHz sampling rate. The master clock (MCLK) for the ADC and DAC is derived from the APLL and set to 256× the sample rate (49.152 MHz) to satisfy audio codec requirements. The I2S input is configured in standard stereo mode with 16-bit samples. Direct Memory Access (DMA) transfers data from the ADCto the PSRAMbuffer without CPU intervention, ensuring minimal dropouts [22]. The output I2S stream is synchronised to the same sample rate, allowing full-duplex operation.

The ESP32-S3 microcontroller is configured to operate in full-duplex I2S mode using separate ports for input and output. The ADC (WM8782) interfaces via I2S0 (input), and the DAC (PCM5102A) interfaces via I2S1 (output). The implementation overview is provided in supplementary material B.4.

### 2.4 Heterodyne Signal Processing and Mix Mode Playback

The Esperdyne firmware implements real-time dual-channel heterodyning on a 192 kHz I2S audio stream. Incoming ultrasonic signals from the stereo ADC (WM8782) are acquired using I2S0 and processed in blocks of 1024 samples. Each channel is heterodyned independently by multiplying the raw audio with a synthetic carrier wave, whose frequency is user-adjustable from 10 kHz to 85 kHz in 5 kHz steps.

Each stereo sample pair’s left and right channels are first normalised to a floating-point range and scaled. These are then multiplied by cosine functions with accumulating phase (phase1 and phase2) calculated as:

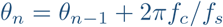

where *f_c_* is the carrier frequency (per channel), and *f_s_* is the 192 kHz sample rate. After phase wrapping and gain adjustment, the output is optionally mixed down to mono if mix mode is enabled. Both channels are averaged and sent to both stereo output channels in mix mode, producing a unified audio stream. If mix mode is off, the heterodyned left and right signals are routed to the corresponding DAC output channels.

To ensure timing coherence, the output is streamed over I2S1 with identical sample rate and bit depth settings, allowing full-duplex operation.

#### Mix Mode Handling

Adedicated toggle button switches between stereo (ST) and mix (MX) modes. The state is reflected on the OLED display and applied in real-time during the audio processing loop. The corresponding logic is conditionally executed per sample block and does not introduce latency beyond buffer processing time.

### 2.5 ADC Configuration

The I^2^S ADC (WM8782) [20] employed in this system is a 24-bit multi-bit sigma–delta converter with integrated digital anti-aliasing filters. In addition to the minimal analogue front-end conditioning, the oversampling modulator and decimation stage provide effective suppression of out-of-band components, with the internal digital filter maintaining a flat passband response up to approximately 0.45 *f_s_* and offering more than 65 dB attenuation above 0.55 *f_s_* [20]. This ensures that signals of interest, including ultrasonic content below the Nyquist frequency, are reliably captured without significant aliasing artefacts. The hardware setup, including I^2^S line configuration, sampling resolution, and master clock synchronisation, follows the design described for the Batsy4-Pro system [15], unless otherwise specified.

Notably, the ADC sampling rate represents the principal limiting factor of the Esperdyne system. Although the ESP32-S3 operates at a core frequency of 240 MHz and includes a dedicated audio Phase-locked loop (PLL) capable of generating master clocks up to approximately 128 MHz [23], the WM8782 constrains acquisition to lower rates. In contrast, the PCM5102A DAC can accept input rates up to 384 kHz, which remain well within the achievable APLL range. Thus, while the processing and clocking capabilities of the ESP32-S3 are sufficient for higher-fidelity acquisition and playback, the present configuration is ultimately limited by the maximum conversion rate of the ADC. Future iterations employing higher-bandwidth converters could therefore exploit the full potential of the system’s clocking and processing resources.

A practical upgrade path for users requiring higher sampling rates is the direct replacement of the WM8782 with a higher-bandwidth I^2^S converter such as the AK5572 (e.g., the KaamosTech AK5572-V2 module) [24]. This converter supports substantially higher sampling frequencies, upto 768kHz, while remaining compatible with the differential MEMS microphones used in the present design. In principle, the hardware modification is limited to substituting the ADCmodule and making minor adjustments to the I^2^S wiring. However, operating at higher acquisition rates substantially increases the data throughput, and the firmware must therefore be adapted accordingly – for example by adjusting ring-buffer sizes and managing PSRAM usage to prevent premature buffer saturation. Preliminary tests conducted on the same ESP32-S3 platform demonstrate stable acquisition at sampling rates up to 250 kHz, indicating that the microcontroller can accommodate higher-bandwidth converters with appropriate firmware optimisation. A fully integrated prototype incorporating this ADC has not yet been field-tested due to seasonal unavailability of bats, but ongoing updates to the open-source repository will include support for higher-fidelity configurations as they are validated.

### 2.6 Power Supply and Switching

The device is powered by a 3.7 V 1000 mAh Li-ion battery connected via a charging module. The module features overcharge protection and provides USB-C recharging support with a standard 5 V DC. The system is turned on and off via a *Single Pole, Double Throw* (SPDT) toggle switch wired to the battery output. Runtime tests showed an operational lifetime of approximately 7 hours on a full charge, depending on output volume and SD card activity.

### 2.7 Data Processing and Curation

To streamline inspection and organisation of field recordings, I developed a custom MATLAB-based graphical interface, *Bat Reviewer* (v1.0), shown in Figure 3. Executable standalone packages for the software are provided for MacOS and Windows operating systems that can be freely downloaded and installed at no cost, with no usage restrictions [18]. The tool provides an integrated environment for browsing, listening to, and exporting ultrasonic recordings collected during field sessions. Files stored in a directory are automatically detected and listed, allowing users to navigate sequentially or by direct selection. This eliminates the need to load each recording manually and supports rapid scanning through large datasets.

The interface’s central panel displays waveform and spectrogram representations of the selected file. The waveform offers a quick overview of the amplitude envelope, while the spectrogram provides a time–frequency view with configurable parameters such as FFT size, window type, and overlap. These options allow the user to adjust the spectral resolution interactively.

For auditory inspection, Bat Reviewer V1.0 incorporates a heterodyne listening function. Users may select any channel in multi-channel recordings and apply a tunable carrier frequency between 15 kHz – (0.5*×F_s_*) kHz. The heterodyned signal is automatically resampled to 44.1 kHz for play-back on standard audio hardware, with an adjustable gain control to optimise listening levels. This functionality enables real-time auditory assessment of call structure, complementing the visual analysis provided by the spectrogram.

Recordings of interest can be copied directly to a user-specified destination folder via the export panel. Aflexible collision-handling policy allows overwriting, automatic renaming, or prompting before replacement, ensuring that curated datasets are built consistently without accidental data loss.

Together, these features allow efficient review and curation of raw recordings, facilitating the identification of high-quality calls and reducing the time required for downstream analysis. Combining visualisation, listening, and file management in a single interface, *Bat Reviewer* provides a practical solution for sorting and preparing large volumes of acoustic data collected in the field.

## 3 RESULTS AND DISCUSSION

The *Esperdyne* prototype was assembled manually on a perforated PCB(Figure 1), with wires hand-soldered between corresponding pins of the components (Figure 2). The fully assembled unit weighed approximately 150 g. To protect exposed solder joints, the underside of the device and contact points were covered with insulating tape.

**Figure 2:**
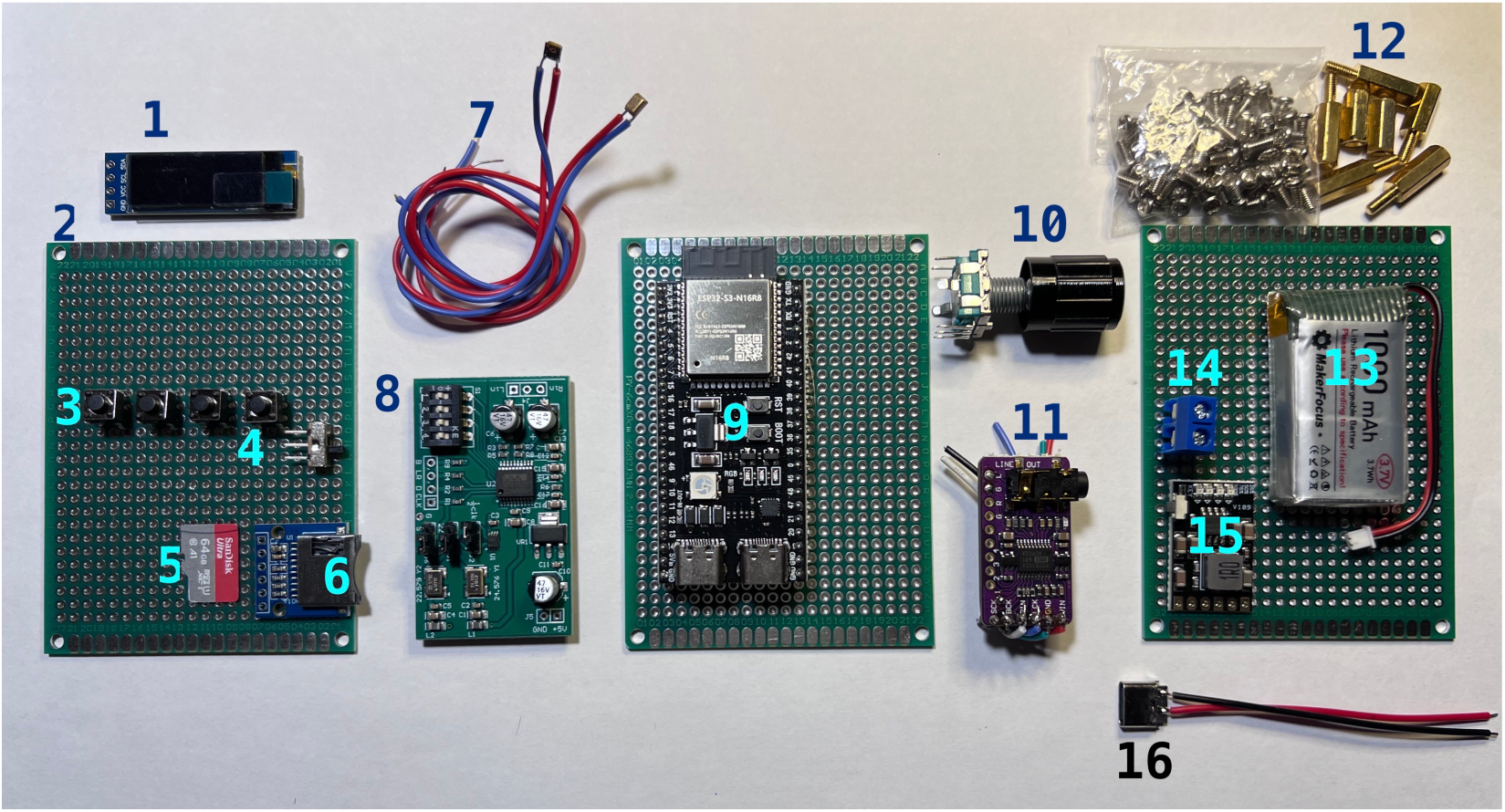
Components of the Esperdyne device: (1) 128×32 OLED Display, **(2)** Perforated PCB **(3)** Tactile push buttons (for control), **(4)** Power On/Off Switch **(5)** microSD card, **(6)** SD Card Module, **(7)** Knowles MEMS Microphones with leads (I2S input), **(8)** WM8782 I2S ADC, **(9)**ESP32-S3 main controller board with USB-C and PSRAM, **(10)**Rotary encoder with push-button, **(11)**PCM5102A stereo DAC breakout board, **(12)** Set of brass hex nuts and screws for mounting, **(13)** LiPo battery (1000 mAh), **(14)** Terminal block for battery input, **(15)** LiPo battery charging module (with protection), **(16)** USB-C Port for battery charging connection. See Appendix A for material cost and sources.

Field testing was conducted along the river Isar, where bat activity was monitored and recorded over 30 minutes. Around 20 recordings were captured as bats foraged in the riparian forest during this time. The files were manually reviewed and imported into a custom application called Bat Reviewer, which allows browsing waveform data from a folder and selectively copying desired files to a target location. This setup enables efficient selection and curation of recordings for further analysis.

A key component of the workflow was the development of the *Bat Reviewer* interface (Figure 3), designed to accelerate the processing and curation of field recordings. While the Esperdyne device provides high-quality ultrasonic data, subsequent analysis requires rapid identification of high-SNR calls among numerous files generated in a session. The GUI addresses this by combining waveform and spectrogram displays with real-time heterodyne playback, adjustable carrier frequency, and volume control. These features allow users to cross-check spectral features visually while listening to heterodyned audio, a process familiar to field ecologists. Files of interest can be copied directly into curated folders using built-in export tools, which handle naming conflicts automatically. This integration of visualisation, auditory assessment, and file management reduces the time required for manual screening and ensures that only the most relevant recordings are retained for further quantitative analysis.

**Figure 3:**
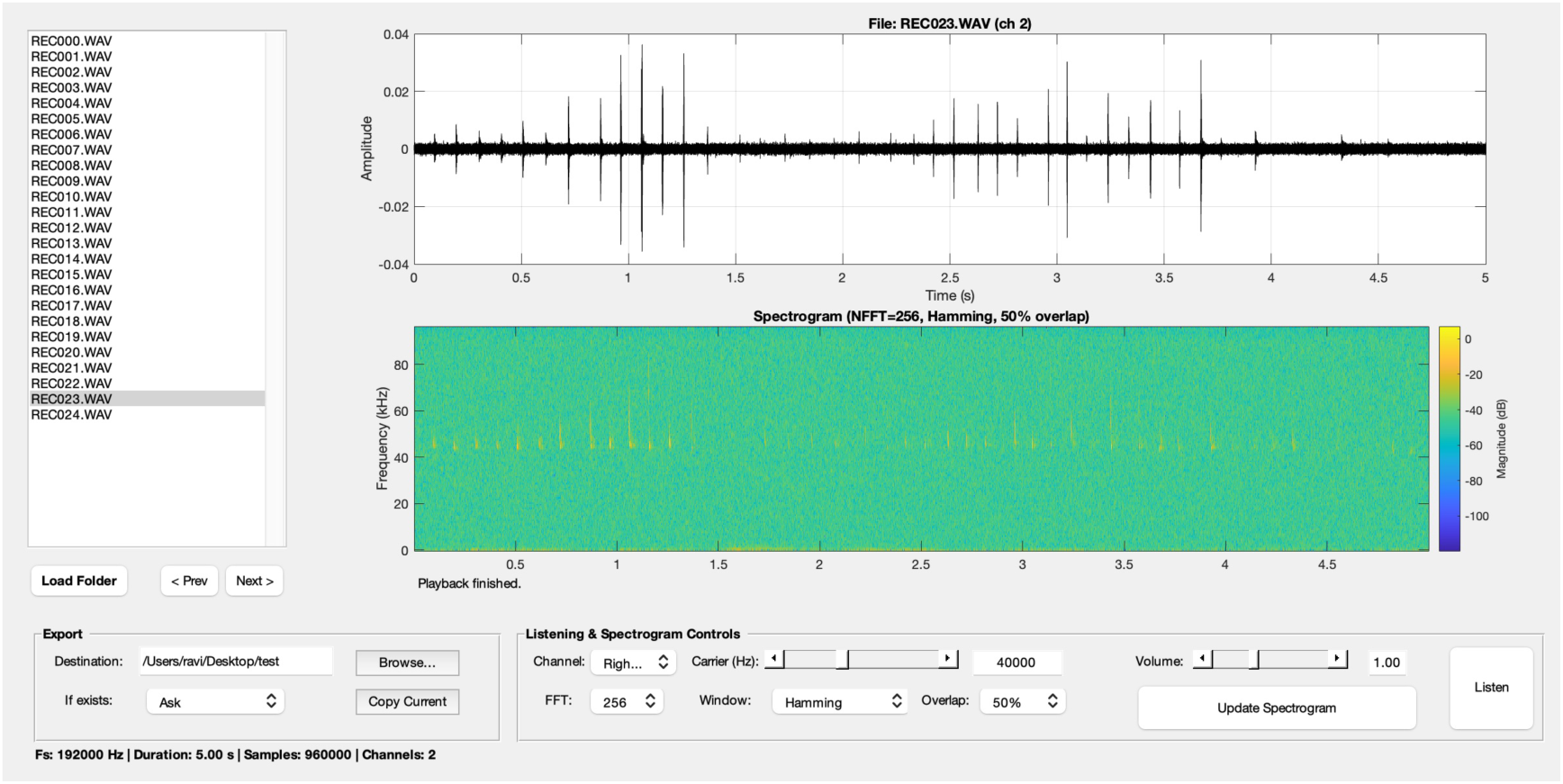
Bat Reviewer V1.0. The interface provides an integrated environment for browsing and visualising ultrasonic recordings. On the left, the file list allows quick navigation through multiple recordings collected in a session, with playback and export options for selected files. The main panel displays waveform (top) and spectrogram (bottom), with user-configurable FFT size, window type, and overlap. Additional controls enable heterodyne listening with adjustable carrier frequency and playback volume. This combination of visual inspection and listening facilitates rapid sorting and selection of recordings of interest from collected datasets.

From my field session, a randomly selected recording was used to extract a sequence of five calls, presented in Figure 4 as both waveforms and spectrograms. As typical of in-field bat recordings, the signals contain both primary calls and multiple echoes—likely arising from ground and vegetation reflections. Notably, the third harmonic was prominently visible as the bat passed the recording site, suggesting a relatively close approach. Despite this, the signal did not clip, owing to the large dynamic range of the selected MEMS microphone.

**Figure 4:**
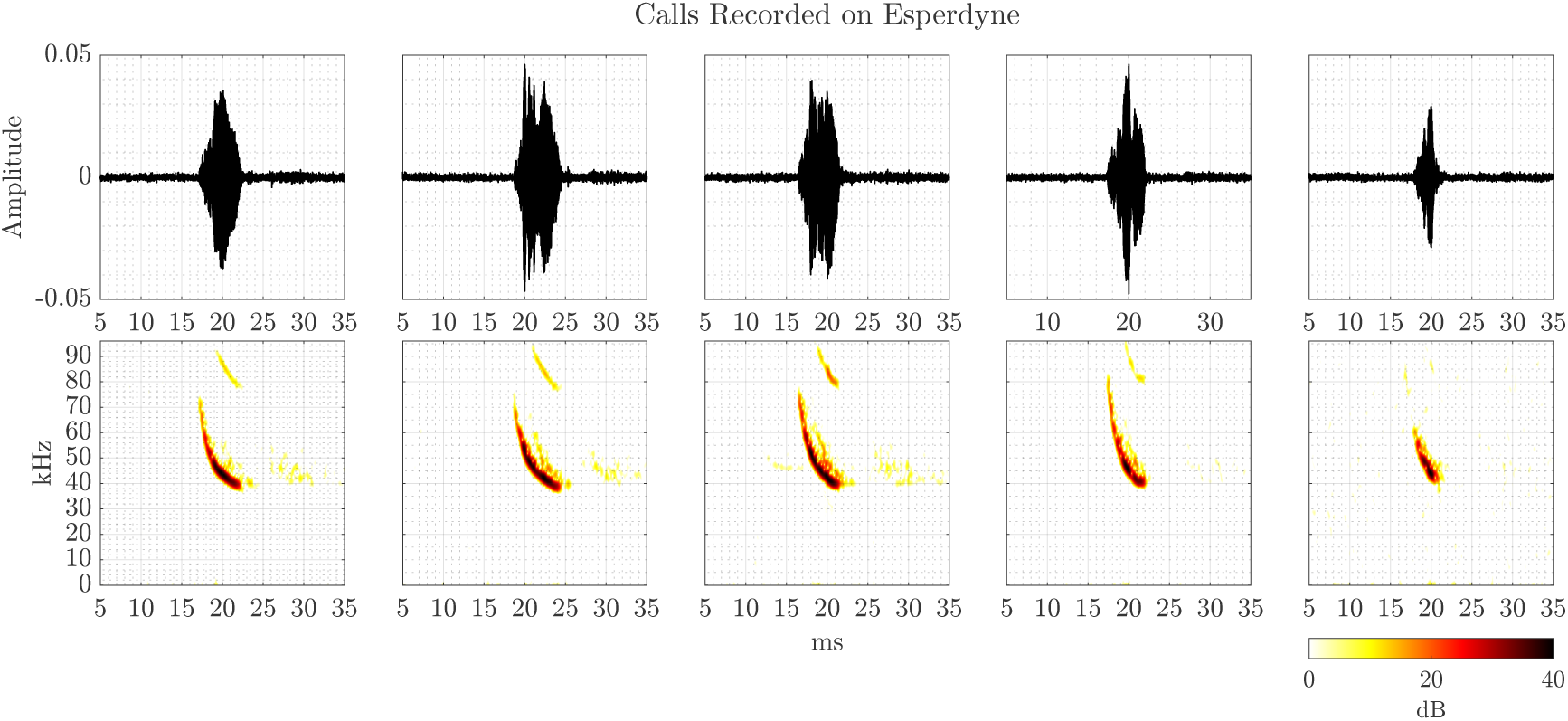
Field-recorded bat call sequence captured using the Esperdyne device. The data were collected in late August 2025 along the river Isar, in an area characterised by thick riparian vegetation. The waveform and spectrogram panels illustrate a sequence of echolocation calls emitted as the bat flew across the microphone. Notably, environmental echoes from the ground and vegetation are visible as smeared features following the primary call pulses, and the third harmonic of the call is distinctly captured.

The wide dynamic range of the microphone is a critical selection criterion for the intended application, particularly given the variability in call amplitude among bat species—some being “whispering” bats. In contrast, others vocalise at levels up to 130 dB SPL [25–31]. I recommend including a microphone preamplifier with adjustable gain for more versatile designs to accommodate a broader range of vocalisation intensities.

A feature to toggle the OLED display on and off was deemed beneficial during testing, as the screen’s brightness can be disruptive in low-light conditions. While this feature may be a personal preference, it can be readily deactivated or customised in the firmware by extending the button press duration required to toggle it, thereby preventing accidental deactivation of the visual display. All other functions remain operational while the display is inactive.

The MX and ST hearing modes were designed with multi-species foraging scenarios in mind, such as those observed in tropical field sites. In particular, the second-channel heterodyne mode enables clearer species separation, especially for CF bats. By default, the MX mode mirrors the signal across both channels, but can be toggled to proper stereo monitoring via a simple push button. Since some field researchers use carrier frequency tuning to estimate species identity, being able to set slightly offset heterodyne frequencies on the two channels enables estimation of upper and lower call frequency bounds—especially useful for FM bats. To the best of my knowledge, such a feature is currently not present in existing commercial detectors.

By default, the system records 5-second stereo audio clips, a choice informed by the observation that high-SNR close-range bat passes—most useful for signal analysis—typically last only a few seconds. While more distant calls are still detectable, they often lack sufficient signal quality. This constraint ensures higher-quality data capture and avoids unnecessary accumulation of low-utility recordings. The included file selection tools support rapid sorting of the acquired session data.

The onboard file counter offers a quick visual indicator of recording progress and session productivity, allowing users to gauge their data yield in real time.

Device development included detailed testing of SD card write speeds. Using the SdFat library and a block size of 4096 samples, a sustained write speed of *≈* 1.5 MB/s was achieved. SPI-based SD access in the ESP Arduino core is typically not optimised for high-speed operations [32, 33], as SD cards are usually used for low-bandwidth tasks like logging text or sensor data. In this application, a 3.8 MB audio file is written in about 3 seconds. A high-speed, ultra-grade microSD card is strongly recommended to avoid data loss or file corruption.

The minimalist UI, purpose-built feature set, and specific affordances for real-time monitoring and retroactive recording make this system a unique and accessible tool. Developed on a widely available microcontroller platform, the device extends the capabilities of ESP-based systems beyond conventional audio libraries, which typically limit users to fixed sample rates optimised for consumer audio. In overcoming these limitations—as I previously did for the Teensy 4.1-based *Batsy4-Pro* [15]—this work contributes to the growing ecosystem of open-source, field-ready tools for global bioacoustic research.

Recent years have seen substantial growth in low-cost and open-source acoustic recorders. Platforms such as Solo demonstrated that inexpensive single-board computers could support flexible bioacoustic monitoring [34], while AudioMoth has become a widely adopted full-spectrum logger, offering high sampling rates, low power consumption, and weatherproof autonomous deployment [13, 35]. Other community-driven efforts, including TeensyBat [36], integrate heterodyne, frequency-division, and full-spectrum modes on more capable microcontrollers, though typically with greater assembly complexity. Smartphones paired with ultrasonic frontends provide portable alternatives, but remain constrained by limited native bandwidth and proprietary app ecosystems [37].

Within this landscape, Esperdyne fills a distinct niche. It is a self-contained, hand-held detector that combines dual-channel tunable heterodyne monitoring with ring-buffered full-spectrum recording at 192 kHz, built entirely on open and easily reproducible ESP32-S3 hardware. Compared with devices like AudioMoth, Esperdyne lacks long-term autonomous operation and higher sampling rates, but provides real-time stereo heterodyne feedback – an uncommon feature in low-cost systems and particularly valuable for teaching and rapid field assessments.

High-end commercial detectors (e.g. Pettersson D1000x, Wildlife Acoustics Song Meter and Echo Meter Touch, Elekon Batlogger-Batcorder) remain the benchmark forlow-noise, high-bandwidth, weatherproof monitoring. However, these systems are costly, closed-source, and generally not user-modifiable. Esperdyne is not intended to replace such instruments, but to offer an accessible, extensible alternative where affordability, transparency, and interactive listening are prioritised. It thus serves as a practical complement to both autonomous open-source loggers and proprietary scientific detectors, supporting broader participation in bat monitoring and acoustic education.

As a development prototype, the presented assembly may be a reference for more robust designs. Specifically, a printed circuit board (PCB) could be created to accommodate existing components or consolidate key ICs into a fully integrated, manufacturable system. All components are commercially available with published specifications, and the firmware is openly released for non-commercial applications. Institutional workshops and collaborative efforts across engineering and biology departments may use this foundation to produce reliable, field-ready hardware suitable for all-weather, all-terrain deployment.

## 4 CONCLUSION

*Esperdyne* demonstrates how low-cost, modular, and fully open-source embedded systems can be leveraged to advance bioacoustic research, especially in regions with limited access to commercial instruments. The presented system lowers the technical and financial barriers to ultrasound-based fieldwork by combining real-time heterodyne monitoring, retroactive recording, and a simple user interface on a widely available microcontroller platform. The inclusion of software tools to browse, audit, and organise field recordings adds a layer of usability critical for researchers and educators on the move. As a derivative of the broader *Embedded Ultrasonics* initiative, Esperdyne complements platforms such as Batsy4-Pro, but fills a critical niche for portable, solo-operation, cost-sensitive deployments. This work invites community-driven evolution by openly sharing the hardware design, firmware, and desktop tools—encouraging interdisciplinary collaboration, local fabrication, and field-ready innovation. Ultimately, devices like Esperdyne widen participation in ecological monitoring and conservation science and reflect a larger shift toward democratised, decentralised scientific instrumentation for a more inclusive future in field biology.

## SUMMARY OF REVISIONS

This revised version (V2) incorporates substantial improvements based on reviewer and editor feedback. The main updates are as follows:

- **Improved accessibility:** The companion software is now distributed as stand-alone executables for macOS and Windows, removing the need for a MATLAB installation and improving accessibility for researchers and educators in low-resource settings.
- **ADC upgrade pathway:** Anew paragraph outlines practical options for replacing the WM8782 ADC with higher-bandwidth alternatives such as the AK5572. Preliminary tests demonstrating stable operation at sampling rates of 250 kHz and above are mentioned, together with notes on firmware implications for future versions.
- **Expanded comparative context:** The manuscript now positions Esperdyne within the broader landscape of low-cost and open-source bioacoustic recorders (e.g. AudioMoth, TeensyBat, smartphone-based detectors), clarifying its complementary niche in education, rapid field assessment, and accessible ultrasonic monitoring.
- **General text and structural improvements:** Sections of the methods, discussion, and figure captions have been streamlined for clarity and coherence. The code snippets have been moved to the *Supplementary Information*

These revisions strengthen the manuscript’s clarity, accessibility, and practical relevance, while preserving its core contribution as an open, low-cost tool for ultrasonic bioacoustic monitoring.

## CODE AND DATA AVAILIBILITY

The following GitHub repositories contain the firmware for Esperdyne and the Bat Reviewer tool:

1. https://github.com/raviumadi/Embedded_Ultrasonics/tree/main/Esperdyne
2. https://github.com/raviumadi/Bat-Reviewer.git
3. Umadi, R. (2025). Bat Reviewer: Version 1.0 - With Installation Packages [Computer software]. Zenodo. https://doi.org/10.5281/zenodo.17870453

## ETHICAL STATEMENT

No human or animal participants were involved in the research in a manner requiring specific ethical approval.

## ACKNOWLEDGEMENTS

The author thanks colleagues and supervisor Uwe Firzlaff at TUM for their support and critical feedback while developing the prototypes and this manuscript.

## SUPPLEMENTARY INFORMATION

## A BILL OF MATERIAL

**Table 1:** Bill of Materials for the Esperdyne System. The listed prices are for reference only and may vary depending on your source of purchase. The indicated prices correspond to costs incurred during development between May and August 2025.

| Item | Description | Count | Unit Cost (€) | Source |
| --- | --- | --- | --- | --- |
| ESP32-S3 DevKitC-1-N16R8 | Microcontroller | 1 | 7.99 | Generic Manufacturer |
| WM8782 | I2S ADC | 1 | 14.90 | Audio-phonics.fr |
| Knowles SPU0410LR5H-QB | MEMS Mics | 2 | 0.50 | Digikey.de |
| PC5102A | I2S DAC | 1 | 7.99 | Amazon.de |
| SSD1306 | OLED Display | 1 | 3.25 | AZDelivery.de |
| SD Schield | File operations | 1 | 3.50 | Amazon.de |
| microSD Card | File storage | 1 | 6.95 | Local purchase |
| Push Buttons | User Input | 4 | 0.50 | Amazon.de |
| Rotary Encoder | User Input | 1 | 1.50 | Amazon.de |
| Circuit Switch | Power Supply ON/OFF | 1 | 0.25 | Amazon.de |
| Perforated PCB | System Base | 3 | 1.50 | Amazon.de |
| Terminal Connector | Battery connections | 1 | 0.25 | Amazon.de |
| Battery Charging circuit | Power management | 1 | 1.50 | Amamzon.de |
| Hexnut Screws | PCB Stacking | Assorted | 5.00 | Amazon.de |
| LiPo Battery 3.7V | Power source | 1 | 5.99 | Amazon.de |
| USB C Connector | Charging | 1 | 0.10 | Amazon.de |
| Wiring | 18/22 gauge coloured wires | Assorted | 5.00 | Amazon.de |
| Total |  |  | 72.67 |  |

## B CODE SNIPPETS

### B.1 Signal Flow Configuration

**Listing 1:**
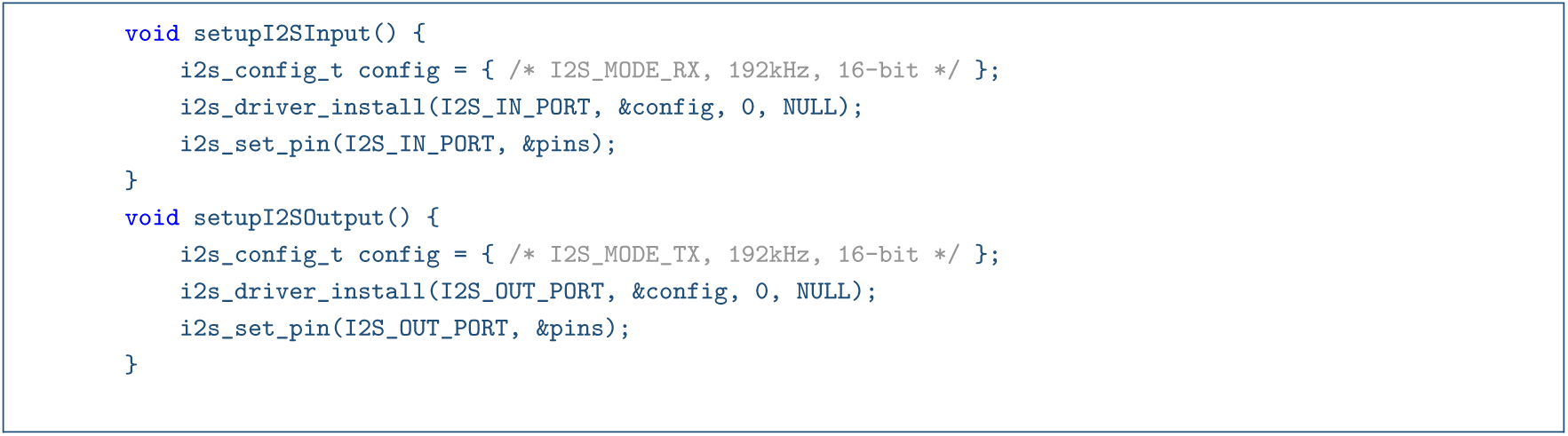
Simplified I2S input/output configuration

### B.2 Ring Buffer Logic

**Listing 2:**
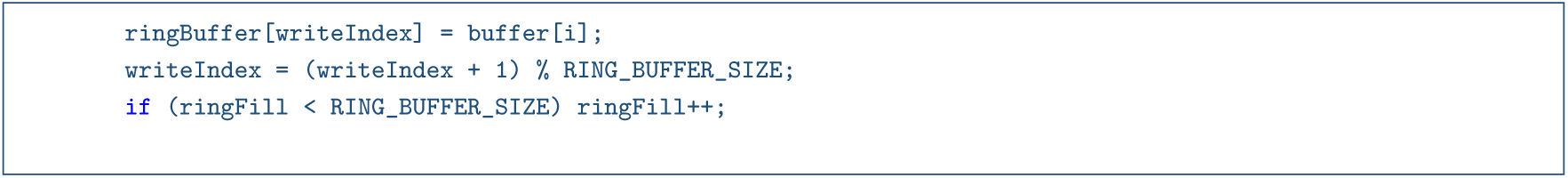
Storing incoming samples to ring buffer

### B.3 User Interaction Logic

**Listing 3:**
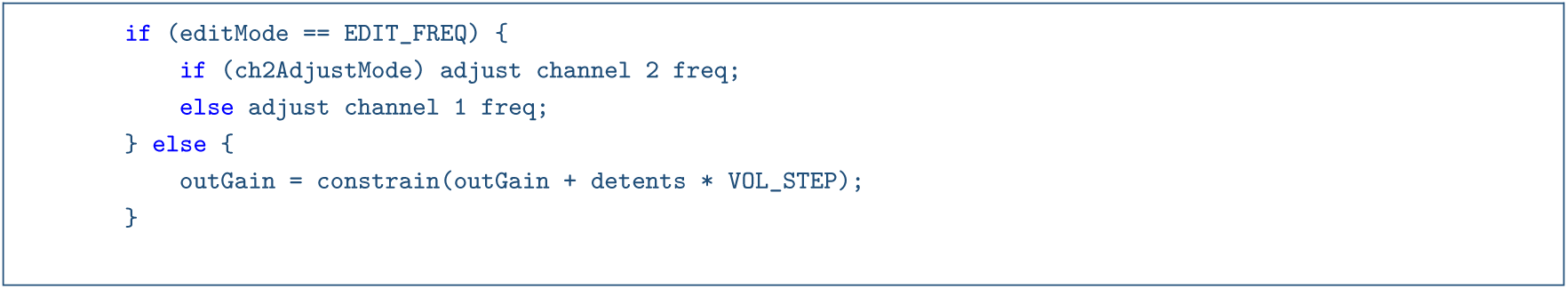
Frequency vs Volume control logic

### B.4 I2S Configuration

**Listing 4:**
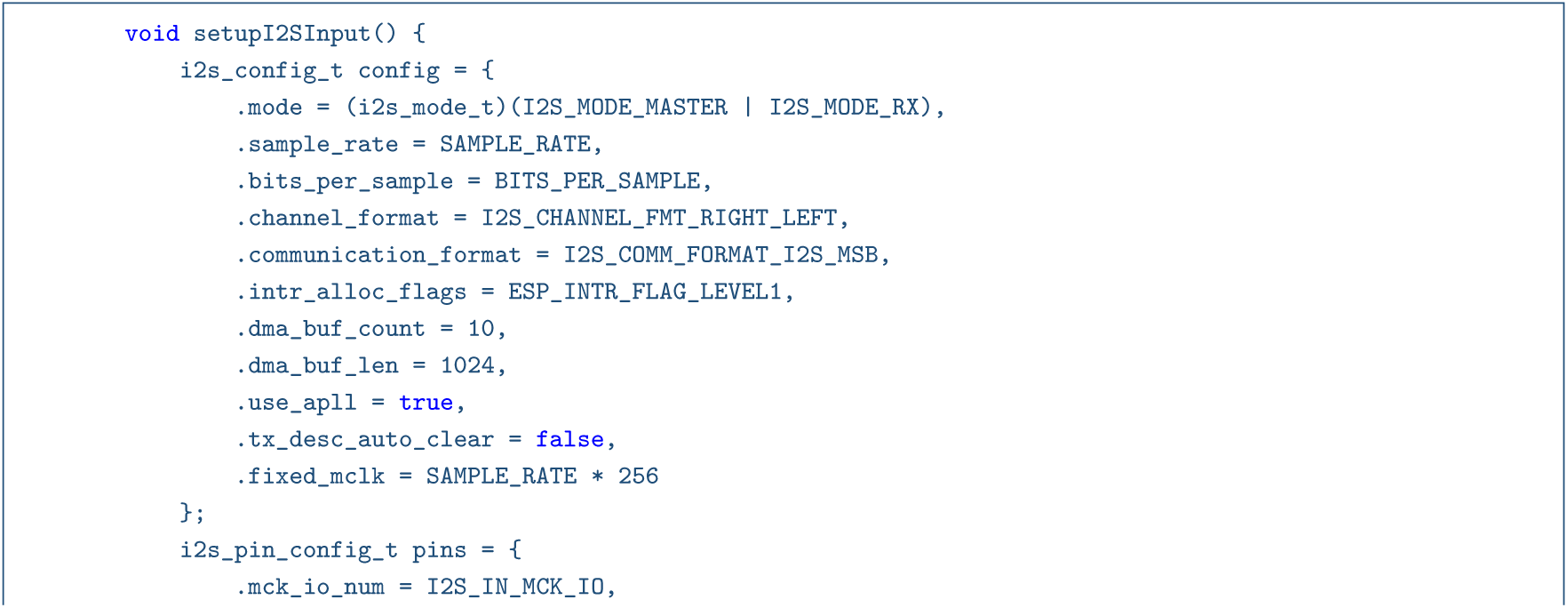
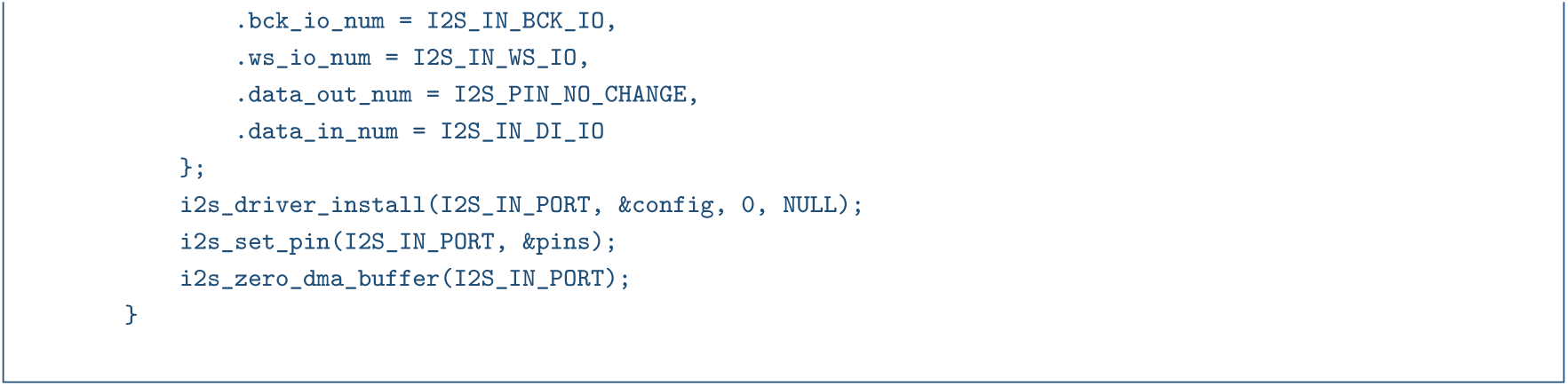
I2S Input Configuration I2S0

**Listing 5:**
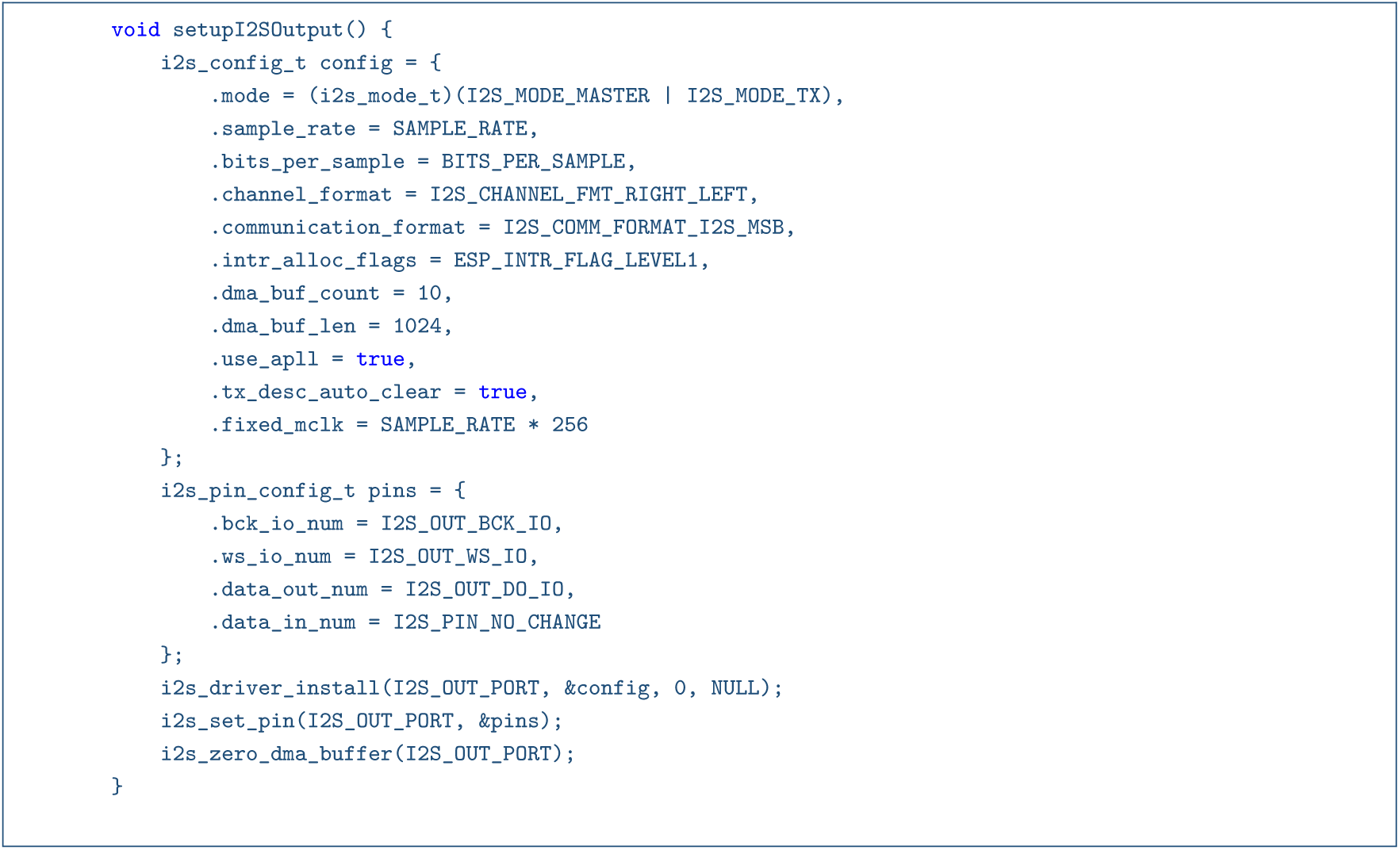
I2S Output Configuration I2S1

## Footnotes

1 The name combines *ESP* (the ESP32 microcontroller) and *dyne* (from heterodyne), with *er* referencing *Vesper*—Latin for “evening” and the root of the bat family name *Vespertilionidae*.

